# Validation of the association between MRI and gene signatures in facioscapulohumeral dystrophy muscle: implications for clinical trial design

**DOI:** 10.1101/2023.02.20.529303

**Authors:** Chao-Jen Wong, Seth D. Friedman, Lauren Snider, Sean R. Bennett, Takako I. Jones, Peter L. Jones, Dennis W.W. Shaw, Silvia S. Blemker, Lara Riem, Olivia DuCharme, Richard J.F.L. Lemmers, Silvère M. van der Maarel, Leo H. Wang, Rabi Tawil, Jeffrey M. Statland, Stephen J. Tapscott

**Affiliations:** Division of Human Biology, Fred Hutchinson Cancer Center, Seattle, WA 98109 USA; Department of Radiology, Seattle Children’s Hospital, Seattle, WA 98105 USA; Department of Pharmacology, University of Nevada, Reno School of Medicine, Reno, NV 89557 USA; Springbok Analytics, Charlottesville, VA 22902 USA; Department of Human Genetics, Leiden University Medical Center, 2333, Leiden, The Netherlands; Department of Neurology, University of Washington, Seattle, WA 98105, USA; Department of Neurology, University of Rochester Medical Center, Rochester, NY 14642, USA; Department of Neurology, University of Kansas Medical Center, Kansas City, KA 66160, USA

## Abstract

Identifying the aberrant expression of DUX4 in skeletal muscle as the cause of facioscapulohumeral dystrophy (FSHD) has led to rational therapeutic development and clinical trials. Several studies support the use of MRI characteristics and the expression of DUX4-regulated genes in muscle biopsies as biomarkers of FSHD disease activity and progression, but reproducibility across studies needs further validation. We performed lower-extremity MRI and muscle biopsies in the mid-portion of the tibialis anterior (TA) muscles bilaterally in FSHD subjects and validated our prior reports of the strong association between MRI characteristics and expression of genes regulated by DUX4 and other gene categories associated with FSHD disease activity. We further show that measurements of normalized fat content in the entire TA muscle strongly predict molecular signatures in the mid-portion of the TA. Together with moderate-to-strong correlations of gene signatures and MRI characteristics between the TA muscles bilaterally, these results suggest a whole muscle model of disease progression and provide a strong basis for inclusion of MRI and molecular biomarkers in clinical trial design.

## INTRODUCTION

Facioscapulohumeral dystrophy (FSHD) is the third most prevalent muscular dystrophy in adults affecting between 1/8,000-1/20,000 individuals world-wide (1). Genetic and molecular biology studies determined that FSHD is caused by the aberrant expression of the transcription factor DUX4 in skeletal muscle (2-4). *DUX4* is a retrogene embedded in the D4Z4 macrosatellite repeat array in the subtelomeric region of chromosome 4 and 10 (5, 6). DUX4 is normally expressed in the early human embryo and regulates a portion of the first wave of zygotic gene activation (ZGA) (7-9). Following its brief expression in the embryo, DUX4 is epigenetically silenced in most somatic tissues including skeletal muscle. Mutations that decrease epigenetic silencing of the D4Z4 repeat result in aberrant DUX4 expression in skeletal muscle where it activates expression of the early embryonic transcriptional program and causes FSHD (10, 11).

Although the facial and upper extremity muscles are often affected first in FSHD, it progresses to involve nearly all skeletal muscles of the body (12, 13). However, unlike some other muscular dystrophies, FSHD is characterized by asymmetric muscle involvement, particularly in the initial phases of disease progression. The asymmetric clinical presentation correlate with changes in muscle MRI characteristics. Many MRI studies support a model of disease progression that initiates with an elevated T2 STIR MRI signal hyperintensity (STIR+), consistent with inflammation or edema, followed by T1 MRI signal hyperintensity resulting from fat infiltration with corresponding loss of the STIR+ signal as fat replaces the muscle tissue (14-20). A recent study evaluated MRI-determined fat infiltration along the whole length of an affected muscle over time and showed that fat infiltration starts at the distal end of the muscle and progresses centrally (21), possibly modifying the model of progression to include early muscle fat infiltration distally before central changes of fat and STIR signals.

Our previous studies (22, 23) showed that the changes in MRI correlate with changes in muscle histopathology, ranging from mild changes of increased extracellular matrix (ECM) deposition and complement activation through stages of inflammation with signs of muscle damage and regeneration, culminating in fat replacement of the muscle. The gene expression signature as measured by high throughput sequencing of RNA from muscle biopsies also showed elevation of gene sets representing ECM, complement, and inflammation that correlated increased disease activity and showed the eventual loss of skeletal muscle gene expression in the later stages associated with fat replacement. It is likely that these progressive changes are caused by the aberrant expression of DUX4 in the muscle because DUX4 regulated genes were expressed at very low levels in the early stages of muscle pathology and progressively increase as the other features of the disease pathology increase. Specifically, using an arbitrary cut-off of DUX4 target gene expression based on RNA sequencing to assign a biopsy as DUX4+ or DUX4-showed a high correlation of STIR+ signal with DUX4+ biopsies (22, 23). A similar study also showed an association between a higher expression of DUX4-target genes and STIR+ muscle and that fat fraction measures increased the sensitivity of identifying muscles with higher levels of DUX4-target gene expression (24).

Our current study was designed to validate the correlation between STIR signal and the DUX4 signature, and to determine whether the disease activity observed in each muscle represented muscle-specific disease progression or a systemic progression with asynchronous manifestation of clinical features, MRI, or muscle markers of disease progression. We enrolled 34 FSHD subjects and performed functional studies, lower extremity MRI, and bilateral biopsy of the tibialis anterior (TA) muscles. Results validate the strong correlation between increased DUX4-signature and STIR+ muscle. Further, our study showed that measurement of fat infiltration across the entire muscle predicted the increased DUX4-signature and conversion to STIR+ status somewhat better than calculations of local fat percentages in the region of the muscle biopsy. Finally, there was a moderate-to-strong correlation between the bilateral TA muscles for both MRI status and the molecular signatures measured by RNA sequencing, particularly the signature consistent with B-cell infiltration. Together, these findings validate our prior study by confirming a strong correlation between MRI status and the DUX4-signature of gene expression and extend the prior study by showing the value of fat infiltration measured across the whole muscle as a predictor of local muscle disease progression. An unanticipated finding of the current study is the correlation of MRI and the gene expression signature between the TA muscles in each subject, particularly the signature of immune cell genes, suggesting a systemic component of disease progression.

## RESULTS

### Study cohort

The study cohort included 34 FSHD subjects, 16 female and 18 male, aged 21 to 69 (mean 47.1±14SD) recruited at three sites (University of Washington Medical Center, University of Rochester Medical Center, and Kansas University Medical Center) of the Seattle Paul D. Wellstone Muscular Dystrophy Specialized Research Center. All participants underwent initial functional testing that included lower extremity dorsiflexor strength measurement, lower extremity MRI, and needle muscle biopsy of both tibialis anterior muscles. Fiducial markers on the mid-portion of the TA muscle were used to identify the MRI characteristics in the region of the muscle biopsy. Muscle biopsies were processed for histology, RNA extraction and sequencing with the composite data shown in Suppl Table 1. DNA was extracted for bisulfite sequencing (BSS) of two regions, one being the *DUX4* exons 2-3 on the distal most repeat unit of 4qA permissive allele(s), and the second being upstream of the *DUX4* coding sequence present in all 4q and 10q D4Z4 repeats. Of the 68 attempted biopsies in the 34 subjects, four biopsy samples (01-0019L, 01-0020R, 01-0029R, 13-0006R) were not adequate for histopathology scoring; three biopsies (13-0006R, 13-0008R/L) in two subjects yielded insufficient RNA for sequencing and the RNA from a fourth biopsy (32-0028L) showed degradation (RNA Integrity number (RIN) = 4.0), excluding these four biopsies from further analysis.

### Transcript-based quality assessment of muscle tissue

Although the intent of the needle muscle biopsy is to determine gene expression in muscle tissue, some biopsies do not capture sufficient muscle tissue to be informative, either because of replacement of the muscle by fat, contamination by blood, or biopsy of fibrotic regions. In this study we used sets of genes to estimate the relative representations of blood (HBA1, HBA2, and HBB), fat (FASN, LEP, and SCD), and skeletal muscle (ACTA1, TNNT3, MYH1), each of which was characterized by a score of cumulated scaled transcripts per million 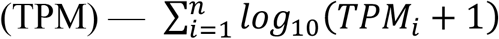. In this bilateral cohort, samples 13-0009R and 13-0007R exhibited low muscle content (below 3% quantile) and elevated fat content (above 97% quantile) (Suppl Fig S1a). The principal component analysis (Suppl Fig S1b) also indicated low-muscle characteristics of these two samples: the loading variables of the first two components revealed low expression in the muscle genes of 13-0009R and 13-0007R. It is reasonable to suspect that these two muscle-low and fat-high content biopsies did not accurately reflect skeletal muscle gene expression. As a result, we excluded them from the analyses of the association of RNA expression levels (e.g., DUX4 signature) to MRI characteristics and other clinical measures.

Based on data from our prior longitudinal study (22) and the current study, we propose applying assessment of muscle, fat, and blood content as a criteria to limit analyses to biopsies that have an insufficient representation of skeletal muscle (Suppl Fig S1c); e.g., identifying samples with a muscle score (cumulated scaled TPM) of less than six or excessive fat score of more than eight as having inadequate muscle tissue for analysis of certain gene categories like DUX4-target genes or the other gene baskets described below.

### Baskets of DUX4-regulated genes distinguish mild-moderate FSHD from control muscle

Yao et al. (25) identified 67 DUX4-regulated genes significantly up-regulated in DUX4-target-positive FSHD biopsy samples, FSHD myotubes, and DUX4-transduced muscle cells. The RNA expression level of this combined set of 67 genes or a subset of four genes (LEUTX, PRAMEF2, KHDC1L, TRIM43) distinguished most FSHD samples from controls and were proposed as candidate FSHD molecular biomarkers. Our subsequent studies determined that mildly or moderately affected FSHD muscles (based on normal MRI and near-normal pathology scores) had modest elevations of these DUX4-regulated genes (22, 23). Therefore, we used these prior datasets to identify subsets of the original 67 DUX4-regulated genes that most reliably distinguished mild-to-moderate FSHD muscle from control. The workflow excluded genes duplicated or deleted in the GRCh38 genome build instead of previously used hg19 (see details in Methods; Suppl Fig 2a). Additional trimming removed the four TRIM49 variants (TRIM49D1/2, TRIM49C, and TRIM49D), whose gene expressions were synchronized with TRIM49; the same applied to three TRIM53 variants and two TRIM43 variants. This process retained 53 of the original 67 genes (Suppl Table 2).

To determine the subset(s) of these 53 genes that best discriminate mild-moderate FSHD muscle from control, we used DESeq2 and Receiver Operator Characteristic curves (with specificity set to 0.8), and also applied a TPM threshold to exclude genes expressed at extremely low levels. This process identified 34 genes as the most robust discriminators for mild-moderate FSHD compared to control muscle (Suppl Table 2). We further refined this to a basket of six (ZSCAN4, CCNA1, PRAMEF5, KHDC1L, MDB3L2, H3Y1) or twelve (PRAMEF15, PRAMEF4, TRIM49, MBD3L3, HNRNPCL2, TRIM43, in addition to the six) of the best discriminating genes. The Random Forest algorithm and leave-one-out cross-validation evaluated the classification performance of all three baskets - with 6, 12, and 34 DUX4-regulated genes. The resulting accuracy rate in classifying mild-to-moderate FSHD samples in the prior longitudinal study data, used to discover these baskets, demonstrated that these baskets outperformed the previously used four genes (LEUTX, PRAMEF2, TRAM43, KHDC1L) (Suppl Fig 2b). Unless otherwise indicated, this study uses the basket of six genes as the DUX4 score (determined by **∑**_***i***_ ***log***_**10**_(***TPM***_***i***_ **+ 1**)).

### MRI measures of regional fat fraction and whole muscle fat infiltration

The local fat fraction at the region of the biopsy ranged from 0.09% to 93.4% (mean 15.7%±25.8%SD) in the 68 TA MRI images; most TAs (54/68) had a regional fat fraction of less than 10%. An alternative method used an artificial intelligence-based approach to generate whole muscle measures of normalized fat content for each TA muscle, termed whole muscle fat percent to distinguish it from regional fat fraction for clarity in this study (see Methods). TA whole muscle fat percent ranged from 3.3% to 82.2% (mean 23.5%± 24.6SD). These two measures, regional fat fraction and whole muscle fat percent, showed strong correlation in cases where regional fat fraction was greater than 10% (Pearson=0.89; Fig. 1a), which corresponded to a whole muscle fat percent of ∼40%. In contrast, muscles with a regional fat fraction below 10% were not well discriminated, whereas the whole muscle fat percent showed a greater linear discrimination in this group with fat percentages ranging between 0 to 40%. These findings are consistent with a prior study showing that whole muscle fat analysis identifies initial changes in fat content at the distal muscle ends prior to the central region of the muscle (21). Indeed, with the exception of muscles that have progressed to very high fat infiltration and lost their STIR+ signal, the STIR-muscles showed distal and, to a lesser extent, proximal fat infiltration prior to central fat replacement; whereas central fat showed greater increase in the STIR+ muscles (Fig 1b). The RNA-seq fat content (based on transcripts of the three fat marker genes, see above) was moderately correlated to the regional fat fraction (Pearson=0.65) and slightly better correlated with the whole muscle fat percent (Pearson=0.69) (Fig 1c). Because of the better discrimination in the low-fat range, we used the whole muscle fat percent measure for the remainder of the analyses.

**Figure 1.**
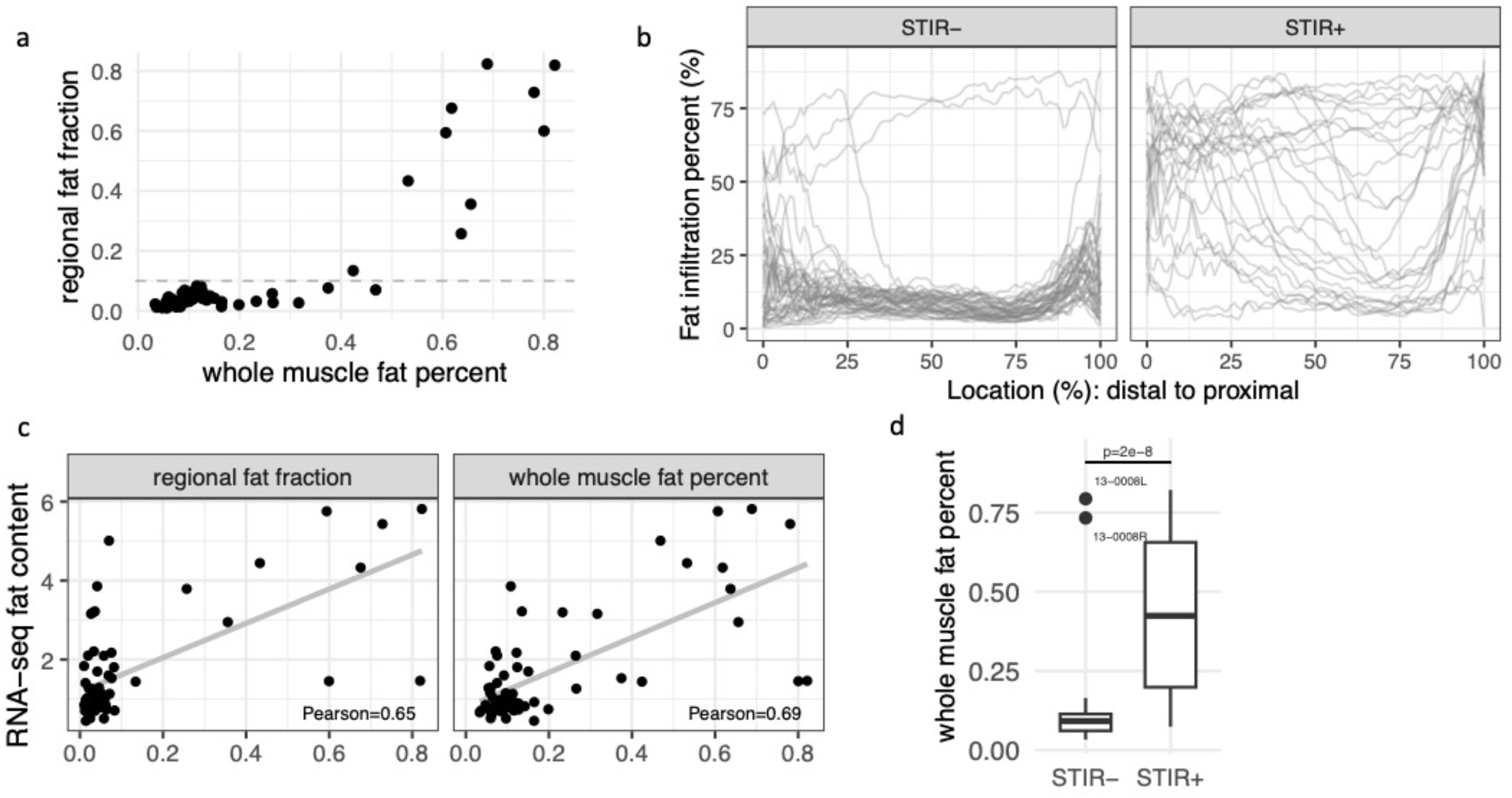
Comparison of regional fat fraction and total muscle fat percentage. (a) At the lower levels of fat infiltration, whole muscle fat percent shows increased values relative to regional fat fraction, possibly due to earlier fat infiltration in the distal and proximal ends of the muscle compared to the region of the biopsy that was generally near the middle of the TA muscle. (b) Fat infiltration percent over the length of each TA muscle from distal (0) to proximal (100) for STIR- and STIR+ muscles. STIR- muscles show fat infiltration in the distal and proximal regions with the exception of muscles with very high fat infiltration that have likely lost prior STIR+ signal, whereas STIR+ muscles show fat infiltration progressing into the central portion of the muscle. (c) Correlation between RNA sequencing inference of fat content with regional or whole muscle MRI measurements of fat content. (d) STIR+ muscles show higher whole muscle fat content with the exception of muscles that have progressed to very high fat percentages.

### STIR, fat infiltration, and DUX4 molecular signature

Of 68 muscles in the 34 subjects, 43 had a STIR rating = 0 (STIR-) and 25 had regions of elevated STIR signal (STIR+). Although STIR signal was rated on a four-point scale (6, 3, 4, and 12 muscles were rated STIR 1, 2, 3, and 4, respectively), our analyses will consider a binary rating of STIR- (rating = 0) or STIR+ (rating = 1-4). STIR+ muscles showed higher levels of whole muscle fat percent (median = 42%) compared to STIR-muscles (median = 9%) (Fig 1d, p=2e-8), however, a subset of muscles with very high whole muscle fat percent (> 73%) were rated STIR-consistent with prior studies showing loss of STIR signal with advanced fat infiltration (note that the biopsies from these two muscles did not yield sufficient RNA for sequencing).

Similar to our prior study showing the association of STIR signal with the DUX4 molecular signature (22, 23), the STIR+ muscles had significantly higher DUX4 scores (mean=3.2) than the STIR-muscles (mean = 0.41), validating a strong association between STIR status and the DUX4 score (Wilcoxon *p*-value = 7e-10). Indeed, muscles with greater than 20% whole muscle fat percent were all STIR+ and all had a relatively high DUX4-score (Fig 2a, DUX4-score plotted on a log scale). The same data plotted with a linear scale for the DUX4-score better shows the decline of the DUX4-score at the higher levels of fat infiltration (Fig 2b).

**Figure 2.**
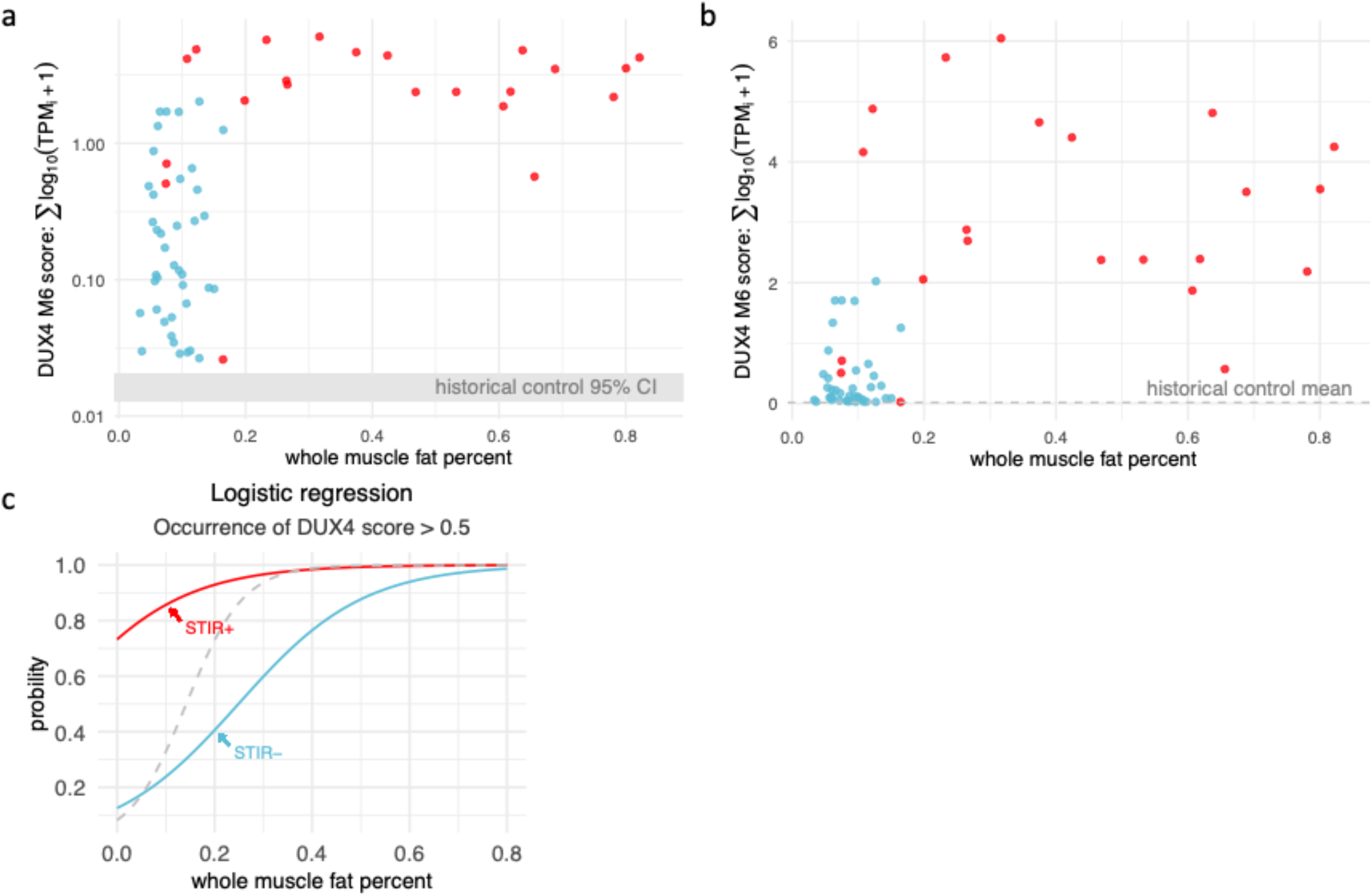
Muscles with whole muscle fat percent greater than 20% and MRI+ muscles have high DUX4 scores. (a) Scatter plot showing that MRI+ muscles mostly have elevated DUX4 scores, and that MRI+ muscles with greater than whole muscle fat infiltration of greater than 20% have uniformly high DUX4 scores. DUX4 scores plotted on a log scale emphasizes the difference between the levels in the historical control muscle biopsies (shaded area indicates the 95% confidence interval (CI) for the distribution of the DUX4 scores in the control biopsies). (b) Same as in (a) but plotted on a linear scale that shows a more bimodal distribution of the DUX4 score, increasing at low-to-mid levels of fat infiltration and decreasing at the higher levels of fat infiltration. (c) Logistic regression predicting the occurrence of a DUX4 score > 0.5. The predictors include the whole muscle fat percent and STIR status of the muscle biopsies and outcome is the occurrence of DUX4 score > 0.5. STIR-muscles indicated in blue, STIR+ in red. Dashed grey line indicates all muscles regardless of STIR status.

The strong correlation between the STIR status and DUX4 score allowed us to build a logistic regression model (see Methods) to predict whether a muscle would have a DUX4-score greater than a specific threshold (Fig. 2c and Suppl Table 3). For example, if the muscle was STIR-with 20% fat infiltration, the probability of the muscle meeting the threshold of a DUX4-score > 0.5 was 41% (Fig 2c, blue line), whereas a STIR+ muscle with 20% whole muscle fat infiltration had a 93% probability of a DUX4-score above this threshold (Fig 2c, red line). Using a very stringent DUX4-score >1.0 yields probabilities of 23% and 70% for STIR- and STIR+, respectively, at the same 20% whole muscle fat infiltration level (Suppl Table 3).

### Association of the DUX4 score to histopathological scores and clinical data

The quantitative myometry muscle strength of the TA ranged from 1.8 to 43 kg (mean 17.2± 10.8SD) and demonstrated an inverse correlation with DUX4 scores (Pearson = -0.5; Fig 3a). The clinical severity score (CSS) showed a moderate positive correlation with the DUX4-score (Pearson = 0.49, Fig 3b). The histopathology score (a rating between 0 to 12 based on variability in fiber size, interstitial fibrosis, central nucleation, and necrosis/regeneration, each ranging from 0-3) also showed a moderate correlation (Pearson = 0.5; Fig. 3c) with the DUX4 score, similar to our prior study (22).

**Figure 3.**
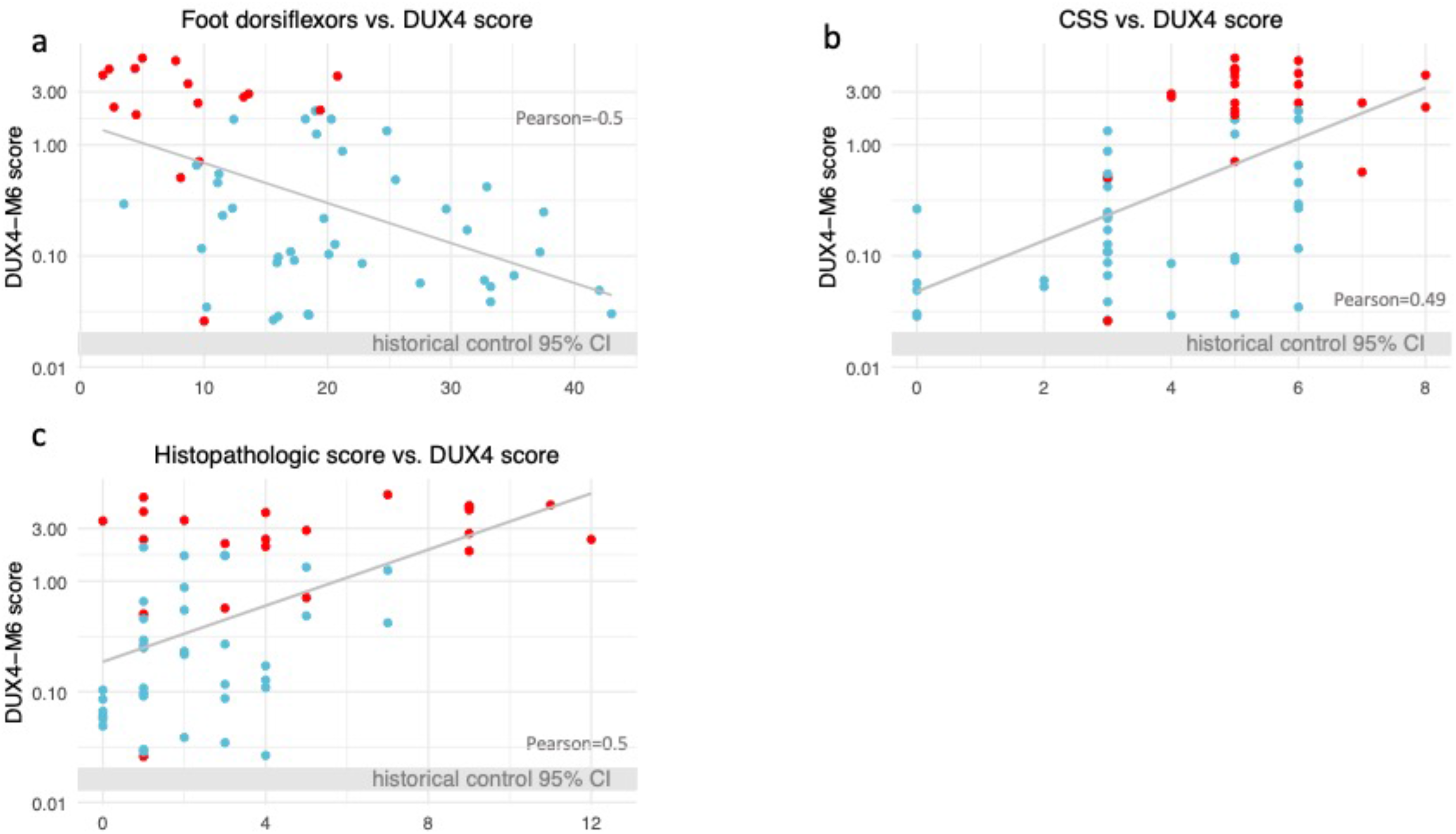
Correlation between functional or histopathological measures and the DUX4 score. (a) Scatter plot showing the correlation between (a) TA muscle strength and the DUX4 score; (b) the clinical severity score (CSS) and the DUX4 score; (c) the histopathology score and the DUX4 score. STIR-muscles indicated in blue, STIR+ in red.

### Inflammatory, ECM, Complement, and Immunoglobulin signatures

In addition to DUX4-regulated genes, our prior studies showed that many other genes had elevated expression in FSHD muscle, including genes related to inflammation, complement, immune cells, and extracellular matrix (ECM) (22, 25, 26). Using a similar approach to the generation of the DUX4-regulated gene baskets, we used the datasets from our prior longitudinal study (22) to identify subsets of genes in each of these categories that most robustly distinguished mild-moderate FSHD from controls (see Methods). By these methods we established baskets of six genes representing signatures for four additional categories: inflammatory (TNC, COL19A1, COMP, THBS1, SFRP2, ADAM12); ECM (PRG4, RUNX1, CCL19, PLA2G2A, CCL18, CDKN1A); immunoglobulin (IGHG1, IGHG2, IGHG3, IGHG4, IGKC, FCGR2B); and complement (C1QA, C1QB, C1QC, C1R, C1S, C3).

Applying these additional gene baskets to the current dataset of TA muscle biopsies validated that the scaled cumulated score (TPM (∑_*i*_ *log*_10_(*TPM*_*i*_ + 1))) of each basket distinguished both STIR- and STIR+ FSHD muscle from the historical control biopsies (Fig 4a) and showed greater elevation in muscles that were STIR+ and had greater than 20% whole muscle fat percent (Suppl Fig 3a and 3b). The expression level of each basket, including the DUX4 basket, was highly correlated with the other baskets across muscle biopsies (Fig 4b). Correlations with other parameters, such as whole muscle fat percent, tibialis anterior strength, sub-categories of histopathologic scores, and CSS are shown in Suppl Fig S3c.

**Figure 4.**
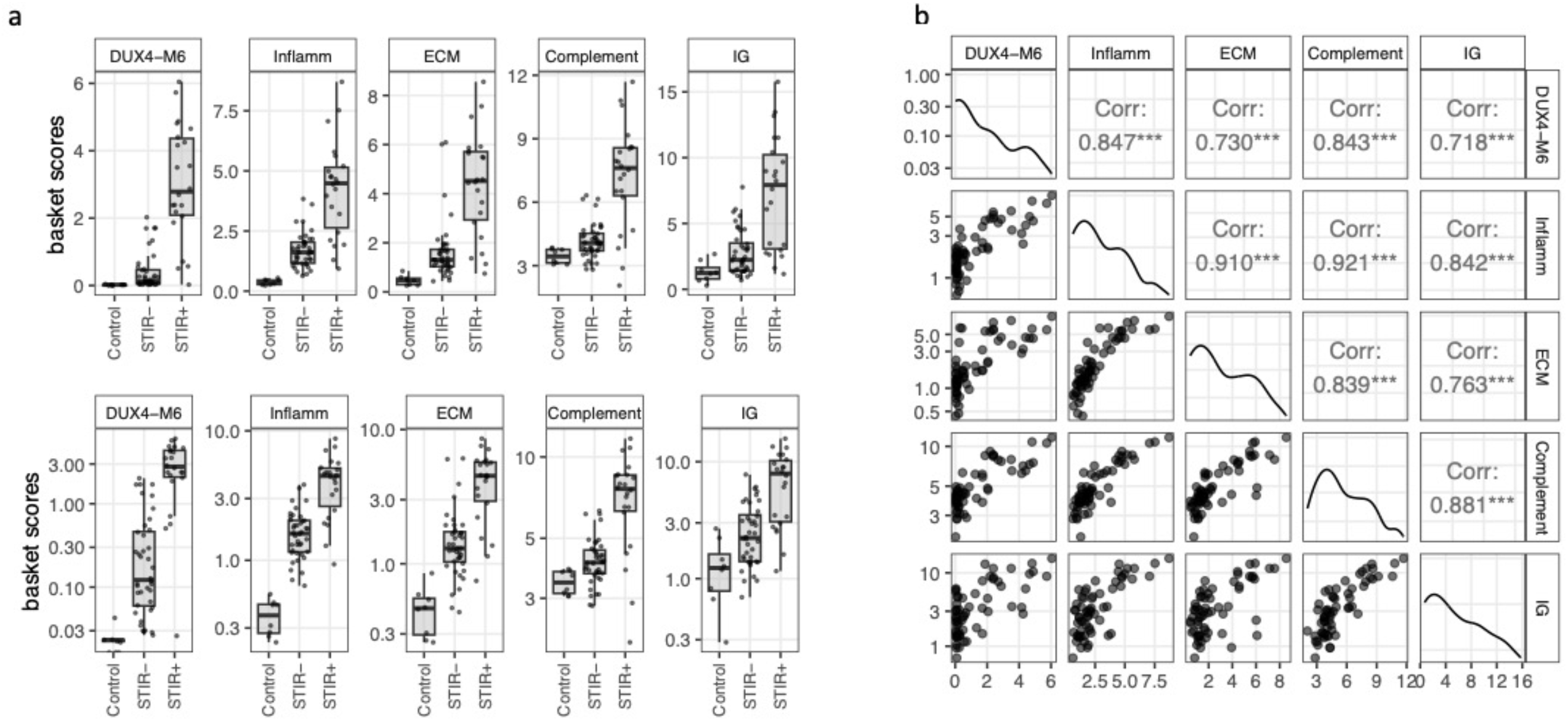
Baskets of genes in different categories distinguish between control, STIR-, and STIR+ muscles. (a) Distribution of the basket scores in FSHD STIR+ and STIR-muscle compared to historic controls. Top panel plotted on a linear scale and bottom panel on a log scale to show separation from control samples. (b) Correlation between the basket scores in each biopsy.

### Bilateral comparisons reveal symmetric trends

Comparisons of the MRI and molecular characteristics of the left and right TA biopsies in the same individual revealed relatively strong correlations. Ten of the subjects were rated STIR+ in both TAs, 19 STIR-bilaterally, and only five discordant for STIR, which differs significantly from the expected values of 4.5 STIR+/+, 13.5 STIR+/-, and 16 STIR-/- (Suppl Fig S4a), determined by random process simulation on pairing 43 STIR+ and 25 STIR-samples (see Methods). Similarly, there is a strong correlation in whole muscle fat percent bilaterally (Pearson=0.82; Fig 5a) and in muscle strength (Pearson=0.94; Fig 5b). Expression of the different baskets of FSHD-elevated genes also showed moderate-to-strong correlation bilaterally with the average Pearson correlations ranging from 0.48 (Inflammation) to 0.82 (IG) (Fig. 5c and Suppl Fig S4b). In contrast, histopathologic scores were not highly correlated (Suppl Fig. S4c).

**Figure 5.**
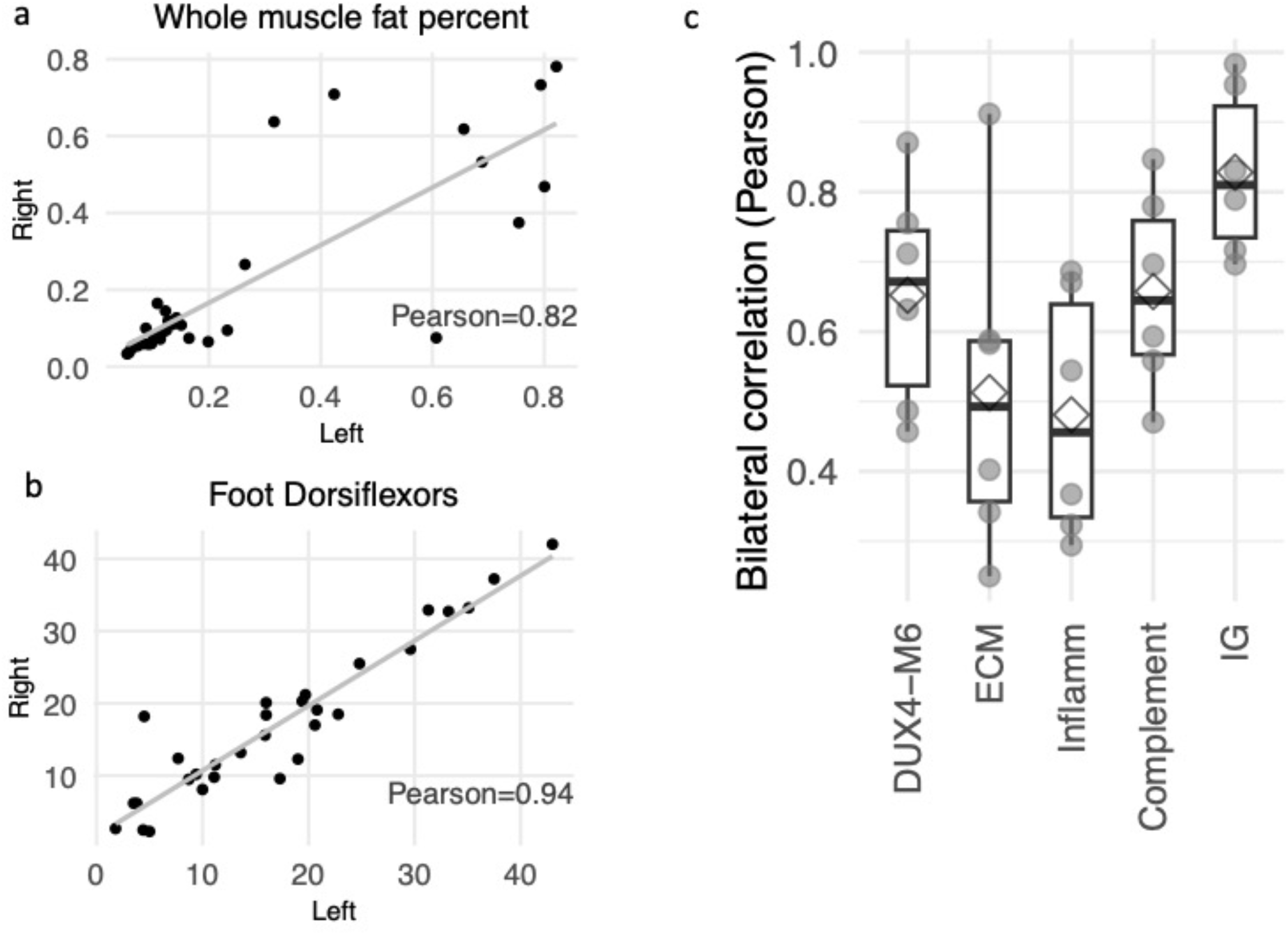
Bilateral comparisons of fat infiltration, TA strength, and gene baskets. Scatter plots show correlation between R and L TA for whole muscle fat infiltration (a) and TA strength (b). (c) Whisker plots showing the correlation between the R and L TA for each basket of genes indicated. Each dot represents the Pearson correlation for each gene in the basket and was calculated based on the gene expression level in TPM. The diamond represents the average of the basket.

### DNA methylation of the FSHD permissive FSHD D4Z4 allele

Of the 68 muscle biopsies collected from 34 subjects, one DNA extraction was inadequate for bisulfite sequencing (13-0006L) and subject 01-0022 had high DNA methylation associated with a duplication of the D4Z4 repeat region and was not included in this analysis. The bisulfite sequencing of subjects with a 4qA161S allele showed a median CpG methylation for the region sequenced of between 0-24%; whereas subjects with a 4qA161L allele had a higher range of 5-43% (mean=16.2%) (Fig 6a and Suppl Table 4). The correlation between the left and right TA biopsies was high (Pearson = 0.85; Fig. 6b). Yet, the methylation levels did not show an association with the STIR status (Fig. 6c) nor demonstrate correlation with other variables such as mRNA levels and clinical scores (Fig. 6d).

**Figure 6.**
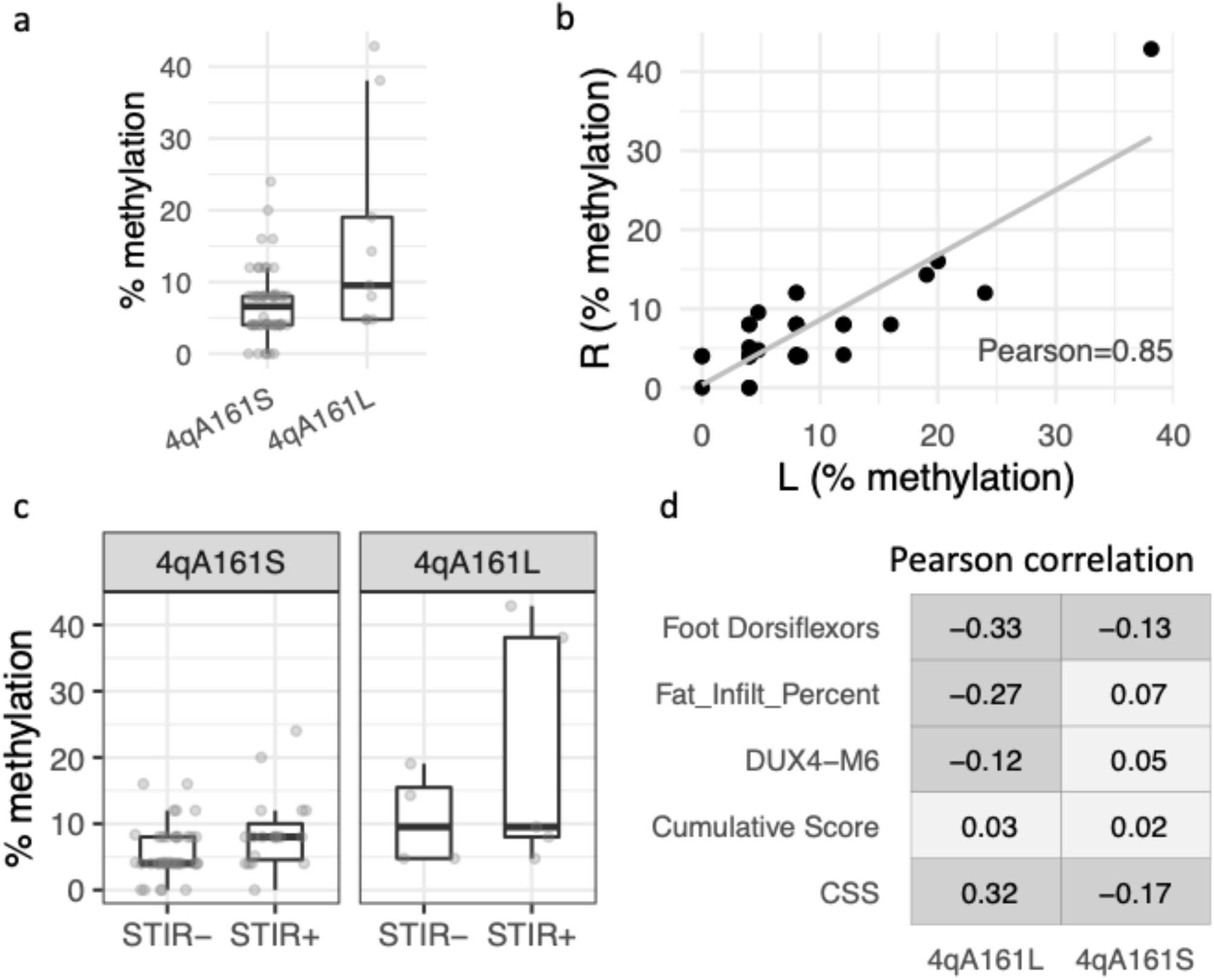
Methylation levels show intra-subject correlation bilaterally not with other parameters. (a) Methylation levels in pathologic 4qA-short and 4qA-long alleles. (b) Bilateral correlation of methylation levels. (c) Methylation levels are not associated with MRI STIR signal. (d) Correlations between methylation levels and the indicated parameter do not show any moderate or strong correlations.

## DISCUSSION

In our previous clinical studies, we used MRI to select a muscle for biopsy and showed a correlation between STIR+ status and exceeding a threshold of DUX4 target gene expression (22, 23). Considerable effort was made to biopsy the region of the muscle with a specific MRI signal characteristic under the assumption that disease progression in the muscle might be focal or regional rather than whole muscle. In the current study, biopsies were performed in the mid-portion of the TA muscles and fiducial targets were used to localize the biopsy region for correlation with regional MRI characteristics. The strong correlation between the molecular signatures and whole muscle MRI characteristics indicates that disease progression is not an entirely focal process, but suggest that the molecular signatures of disease progression spread through the entire muscle even at early stages. This might have significant implications for the design of clinical trials measuring response to therapies because it suggests that a biopsy in a central region of a muscle can accurately reflect the stage of disease progression in that muscle as a whole.

This conclusion is further strengthened by the calculations of whole muscle fat infiltration. Whole muscle fat infiltration showed a stronger association with the molecular signature than the similarly calculated regional muscle fat infiltration in the area of the muscle biopsy. Because fat infiltration begins in the distal regions of the muscle (21) and the biopsy was performed in the central portion of the muscle, this finding also indicates that the molecular signature reflects a whole muscle level of disease activity rather than only focal or regional inflammation or disease. If confirmed by additional studies, this suggests that stringent MRI-designation of a specific biopsy site is not necessary to capture disease activity and will significantly simplify clinical trial design.

In addition, the moderate-to-strong correlations between measurements of disease activity in the bilateral TA muscles also suggests that FSHD progression is not only driven by focal events in an individual skeletal muscle but might also reflect a systemic progression of the disease. Although speculative at this time, the high correlation of immunoglobulin mRNAs in the bilateral TA muscle biopsies raises the possibility that a systemic immune response to FSHD antigens might have a role in coordinating disease progression in different muscles of an individual.

In summary, this study validated the previously identified strong correlation of an elevated DUX4-signature with STIR+ muscles and revealed unanticipated findings that support a model of entire muscle and even systemic disease progression. First, the DUX4-signature at a biopsy site in the mid-portion of the muscle correlates with the entire muscle MRI signal, indicating that progression is not entirely regional or focal. Second, the moderate-to-strong correlations between bilateral TA muscle molecular signatures support a model that incorporates an element of systemic disease progression, perhaps mediated by circulating factors such a B or T cells. Our current study supports using MRI determinations of STIR and whole muscle fat infiltration together with molecular signatures as measures of disease progression in FSHD in future clinical trials.

## MATERIALS AND METHODS

The study was conducted jointly at the University of Washington, the University of Rochester, and University of Kansas through the Seattle Paul D. Wellstone Muscular Dystrophy Cooperative Research Center. The study was approved by the Human Subjects Committee at each institution, with written informed consent obtained for all participants. Patients were examined and given a 10-point Clinical Severity Score (CSS).

### Magnetic Resonance Imaging (MRI)

<u>All</u> MRI examinations were performed on a 3T Siemens scanners running E11C. Fiducial stickers placed at the time of MRI were used for biopsy targeting of the TA’s and localizing muscle features. T1/T2 Dixon and STIR sequences were centered around the tibial spine (upper/lower station) and acquired using flexible array coils. Sequence parameters were as follows: T1_Dixon: TE1=1.35ms, TE2=2.58ms, TR=4.12ms, matrix=448×266, voxel size=1.1×1.1×4.0mm, 104 slices; T2_Dixon (2 echoes): TE=97, TR=5570ms, matrix=448×364, voxel size=1.1×1.1×8.0mm, 40 slices; STIR: TE=38ms, TR=3150ms, matrix=320×220, voxel size=1.1×1.1×5.0, 40 slices).

#### MRI analytics

Data for each subject were converted into slicer and regions-of-interest created to approximate the punctate biopsy regions (∼0.1±.03cm) using fiducial landmarks visible on the scans to generate regional fat estimates and UTE measures. STIR hyperintensity were rated qualitatively (by DS and SF) on a four-point scale: 0, normal appearance; 1, very mild diffuse elevation/may be artifact, close to zero; 2, mild diffuse elevation; 3, moderate areas of increased signal intensity; 4, severe involvement of entire muscle.

T1-weighted Dixon scans were processed using a combination of an artificially intelligent (AI)-based algorithm and manual vetting (Springbok Analytics, Charlottesville, VA). In brief, the AI algorithm segmented the TA from all MRI axial slices (27). The AI-based segmentation was then vetted by a team of trained segmentation engineers and evaluated by a single researcher (OD) to ensure consistency of segmentation and provide a final 3D label of the TA. The TA volume was calculated by summing the voxel volumes from the segmented TA labels on all slices. To reduce the effects of body size on muscle size across patients, muscle volume was normalized by the product of the patient’s height and mass. Lower extremity muscle volumes have been previously shown to vary with the product of height and body mass in healthy, active subjects (28). To minimize the effect of varying coverage ranges on total TA volume captured; the TA volume was further normalized by the anatomical muscle length of the TA. The anatomical length for the TA was calculated by taking the sum of the 3D Euclidean distances between adjacent centroids of the TA in axial slices (28). Fat infiltration % was found for each pixel and was calculated as the pixel’s fat series intensity divided by the sum of the pixel’s fat series intensity and water series intensity (29). Whole TA fat infiltration was found by taking the average fat infiltration across all pixels corresponding to the labeled TA. A vectorized version of fat infiltration along muscle length was found by computing fat infiltration of the TA at each axial slice as a function of axial slice progression along the TA (from distal to proximal). As described in the text, values of fat within the regional sample, entire muscle, and STIR features were reduced for comparison to RNA data.

### Muscle Biopsy for Pathologic Grading and Biomarker Studies

Subjects underwent bilateral tibialis anterior muscle biopsy. Fiducial targets on TA were used to localized confluent STIR+ regions in the mid-portion of the TA if present. Biopsies were obtained under sterile condition using either UCH modified or unmodified Bergstrom needles.

### Muscle pathology grading

The histopathologic samples were graded for the severity of the pathologic changes based on 10 µm sections stained with Hematoxylin & Eosin and Trichrome. A pathologic severity score is determined for each biopsy based on a 12-point scale giving a 0-3 score to each of four major histologic features: variability in fiber size, percent of centrally located nuclei, interstitial fibrosis, and muscle fiber necrosis/regeneration/inflammation.

### RNA-seq library preparation and sequencing

RNA was extracted from frozen crushed muscle biopsy material using TRIzol and single end 100 nt Illumina sequencing was performed by the Fred Hutchinson Cancer Center Genomics Core.

### RNA-seq preprocessing and data analysis

The RNA-seq preprocessing pipeline started with filtering out unqualified raw reads and trimming the Illumina universal adapters by Trimmomatic, followed by alignment against genome built GRCh38(p13) by Rsubread. The gene features were collected from GENCODE, version 35 and the gene counts were profiled by using the GenomicAlignment::summarizedOverlap() function with “IntersectionStrict” mode, counting reads that completely fall within the range of exons and ignoring ambiguous reads straddling different gene features. The normalization, regularized log transformation and differential analysis were performed by DESeq2.

### Curation for baskets of DUX4 and other FSHD-specific signatures

Here, our goal is to curate a subset of DUX4-targeted genes to represent the DUX4 signature with better performance in discriminating mild-to-moderately affected muscles on molecular levels. We started with 67 DUX4-targeted genes that are robustly up-regulated in DUX4-target-positive FSHD biopsy samples, FSHD myotubes, and DUX4-transduced muscle cells (25), followed by

1. lifting the coordinates and annotation from the initially used hg19 to GRCh38 genome build,
2. removing duplicated or deleted DUX4 targeted genes, and
3. removing four variants with synchronized expression of TRIM49, three of TRIM 53, and two of TRIM43.

To determine the subset(s) of the retained genes that best discriminate the mild-modest FSHD muscles from control, we performed a differential analysis comparing the previously classified Mild and Moderate RNA-seq samples from the prior longitudinal study (22) to the historical controls (23). The approaches include (1) the classification-based Receiver Operator Characteristic curves (ROC) with a specificity of 0.8 and (2) the hypothesis-based DESeq2 with the significance criteria set to adjusted p-value < 0.05 corresponding to ℋ_0_ *log*_2_|*FSHD*/*control*| < 1. We further excluded genes expressed at extremely low levels by setting a TPM threshold—average TPM > 2 on the RNA-seq samples classified as the High and IG-High classes. Consequently, we identified 34 genes as the most robust discriminators for the mild-moderate FSHD biopsied muscles compared to the controls (Suppl Table 2). The further refined baskets of the top-ranked genes (by DESeq2 and ROC) include a basket of six (ZSCAN4, CCNA1, PRAMEF5, KHDC1L, MDB3L2, H3Y1) and twelve (PRAMEF15, PRAMEF4, TRIM49, MBD3L3, HNRNPCL2, TRIM43, in addition to the six).

Next, we curated the inflammatory and extracellular matrix basket genes similar to the DUX4 basket. We started with the up-regulated genes (in the mild-modest affected biopsies) associated with inflammatory response and extracellular matrix and ranked them by DESeq2 and ROC. In addition, we selected the top-ranked and best at showing stability between the initial and follow-up visits based on Pearson correlation. As a result, we curated TNC, COL19A1, COMP, THBS1, SFRP2, ADAM12 for the inflammatory basket and PRG4, RUNX1, CCL19, PLA2G2A, CCL18, CDKN1A for ECM.

Finally, for the complement pathway and immunoglobulin baskets, the curation started with the genes associated with complement classical/alternative activation pathway (GO:0006956, GO:0006957, and GO:0006958) and humoral immune response mediated by circulating immunoglobulin (GO:0002455). Since only a few of these genes were differentially expressed in the mild-modest affected muscle, we selected the genes that are up-regulated (by DESeq2, adjusted p-value < 0.05) in more affected biopsied muscles (previously classified as IG-High and High classed). The final list preferred the more expressed and stable between the left and right TA muscles (by Pearson correlation). Hence, we obtained IGHG1, IGHG2, IGHG3, IGHG4, IGKC, and FCGR2B for the immunoglobulin basket and C1QA, C1QB, C1QC, C1R, C1S, C3 for the complement activation basket.

### Classification using FSHD molecular signatures and machine learning

Based on the gene expression profiling, we aimed to classify the TA biopsied muscles as Mild and Moderate+ classes using random forest and FSHD molecular signatures, represented by the DUX4, ECM, Inflammatory, Complement, and IG basket genes. To build the training mode, we employed the RNA-seq profiling from our longitudinal study (22) as the training set where the FSHD biopsies were previously classified (by k-means) into Mild, Moderate, IG-High, and High classes. (Note that the k-means clustered the Mild class with the control samples.) The training metric included two sets of samples: Control and Moderate+, composed of the Moderate, IG-High, and High classes. The leave-one-out cross-validation on the random forest training model yielded 90% accuracy in distinguishing the Moderate+ from the controls. We thus applied the training model to the bilateral TA biopsies, excluding two muscle-low samples, and resulted in 17 Mild (control-like) and 45 Moderate+ samples.

### Logistic regression to predict the dichotomy outcomes using MRI characteristics

Using generalized linear models (GLM), we confirmed that the DUX4+/- dichotomous status of the biopsied muscles (partitioned by the threshold of DUX4 score at 1) is linearly associated with the STIR+/- status (p=1e-5) and whole muscle fat percent (p=3e-3). This linear relationship hinted that we could use a logistic regression model with MRI variables as predictors to predict the odds of DUX+/- outcomes of a TA muscle. To test the stability of the prediction, we performed the same method on our longitudinal and bilateral TA studies using the STIR+/- and regional fat fraction as predictors. The predictions of the two studies yielded similar results (Suppl Fig S3).

### Simulation on pairing samples from two pools of STIR status

We performed 1000 runs of simulation, in which each run, 34 pairs were randomly drawn from 43 STIR- and 25 STIR-samples, yielding the numbers of STIR+/+, STIR-/-, and discordance STIR+/- pairs. This simulation built three distributions of the number of three types of pairs, and the expected values of each pair type are 4.5 pairs for STIR+/+, 13.5 for STIR-/- and 16 and STIR+/- (Suppl Fig S5a).

### DNA methylation analysis

The Trizol/chloroform extracts were frozen after RNA isolation and used for the isolation of genomic DNA from the same biopsy samples. The SSLP haplotype was analyzed as described (https://www.urmc.rochester.edu/neurology/fshd-center/research-info/protocols.aspx#4) on genomic DNA samples to confirm their identities. Genomic DNA (0.6-1.5µg) was bisulfite (BS)-converted and processed per manufactured instructions (EpiTect Bisulfite kit, Qiagen). The BS-converted genomic DNA was used for bisulfite sequencing (BSS) analysis using the 4qA (300ng), 4qA-L (150ng) and 4qA DUX4 5’ (150ng) amplicons, as described in (30, 31), with the following modifications: BS-PCR products were sequenced by next-generation sequencing (NGS) using the Ion Chef and Ion GeneStudio S5 System (Thermo Fisher Scientific).To accommodate the NGS, the forward and reverse BS-PCR primers used in the nested BS-PCR were fused with Ion Xpress barcode adaptor per manufacturer’s instructions (Ion Amplicon Library Preparation User Guide, PN 4468326 Rev C). Sequences were analyzed to provide >3000 independent nonidentical sequencing reads per amplicon.

## Code and data availability

The raw RNA-seq read files will be deposited to Gene Expression Omnibus with accession numbers GSExxxx (pending). A collection of processed data—gene counts, annotation, clinical scores and meta data—are available in R/Bioconductor formatted datasets from our GitHub repository: https://github.com/FredHutch/Wellstone_BiLateral_Biopsy. The repository also hosts a GitHub-page Book, with detailed narratives of analysis and reproducible R codes (https://fredhutch.github.io/Wellstone_BiLateral_Biopsy).

## Software

The statistics analysis and visualization were performed using R/4.2.2, Tidyverse packages, and Bioconductor v3.16 packages; the most frequently used packages are tibble, dplyr, ggplot2, DESeq2, ROC, caret, GenomicAlignments and bookdown. Bioinformatics tools used for preprocessing data include fastqc, Trimmomatic (0.32), SAMtools (1.10) and Rsubreads (2.8.1).

Tabular data from MRI-derived measures are available on reasonable request.

## Acknowledgements

This work was supported by National Institutes of Health [P50 AR065139 to SJT]; and the Friends of FSH Research. The authors thank the FSHD community for their extraordinary help and participation in the study despite the difficulties imposed by the COVID pandemic.

## FIGURE LEGENDS

**Supplemental Figure S1.**
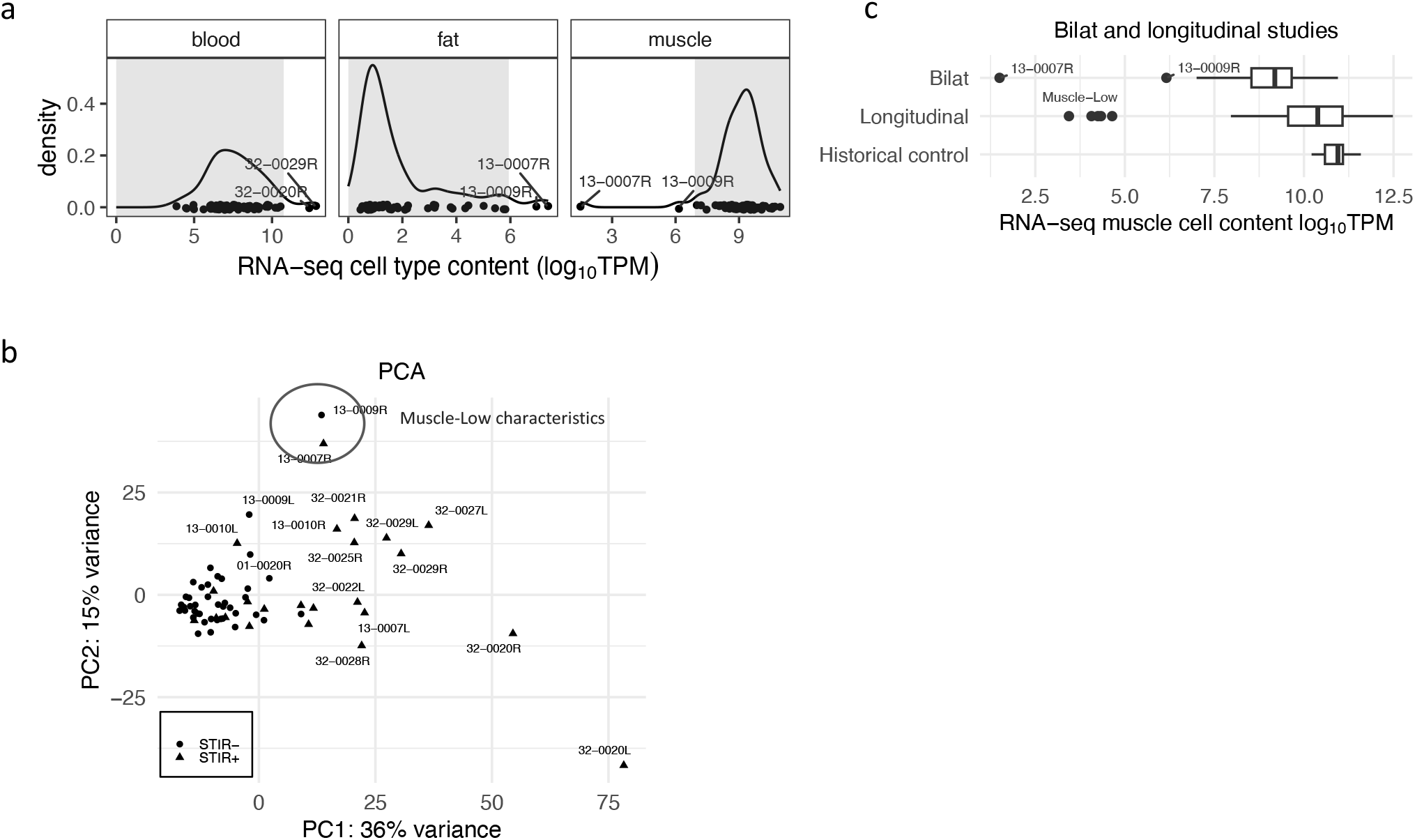
Assessing blood, fat, and muscle RNA to infer representation of muscle and other cell types. (a) Distribution of the cell type content expression: blood cell type characterized by HBA1, HBA2, and HBB; fat by FASN, LEP, and SCD; muscle by ACTA1, TNNT3, MYH1. Gray areas indicate 0-97% quantile for the blood and fat types and 3%-100% for muscle. (b) Principal component analysis on the bilateral biopsies, presented by the first and second principal components, indicates that 13-009R and 13-007R demonstrate low muscle characteristics: low muscle content and elevated expression in fat and immune infiltration. (c) Muscle RNA content from the historical control, prior longitudinal study and the current bilateral study. Note that the “Muscle-Low” group from the longitudinal study was classified by k-mean clustering algorithm as it demonstrates the low-muscle characteristics. In addition, this group has low muscle content scores (2.5 – 5), which indicates that it is reasonable to group 13-0007R and 13-0009R of low 3% quantile muscle content to be “muscle-low”.

**Supplemental Figure S2(a).**
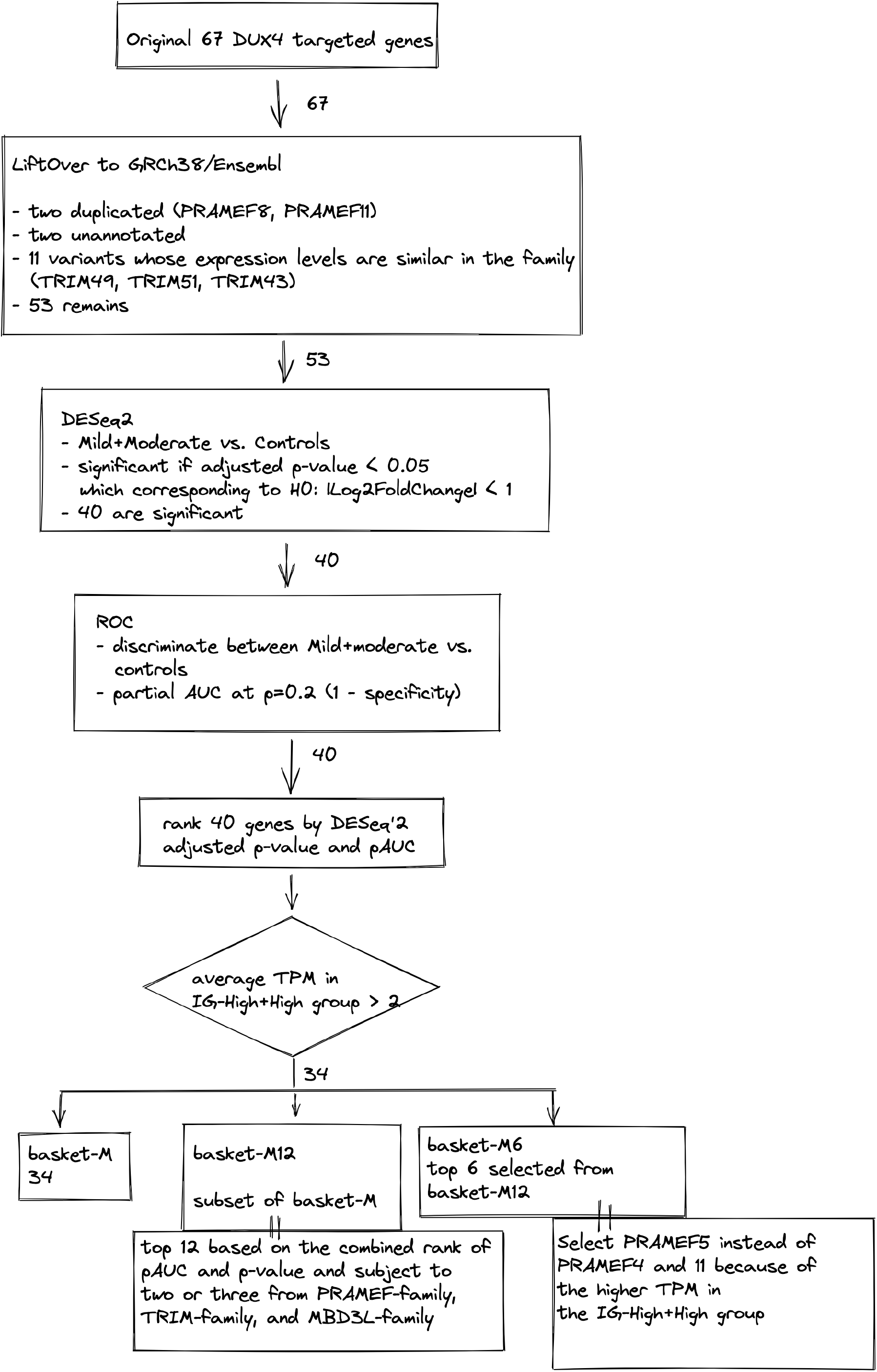
Flowchart of curating baskets of DUX4 biomarkers – DUX4-M6, DUX4-M12, and DUX4-M34. We lifted the original genome reference hg19 to GRCh38 and excluded the unannotated, duplicated, and variants with similar expression levels in the family (keeping one from each family of TRIM49, TRIM51, and TRIM43). We then used DESeq2 and ROC to curate the DUX4 biomarkers that best discriminate the mild + moderate biopsies from the controls in the prior longitudinal study dataset. Along with other criteria listed below, the resulting baskets obtain six, twelve, and 34 genes.

**Supplemental Figure S2(b).**
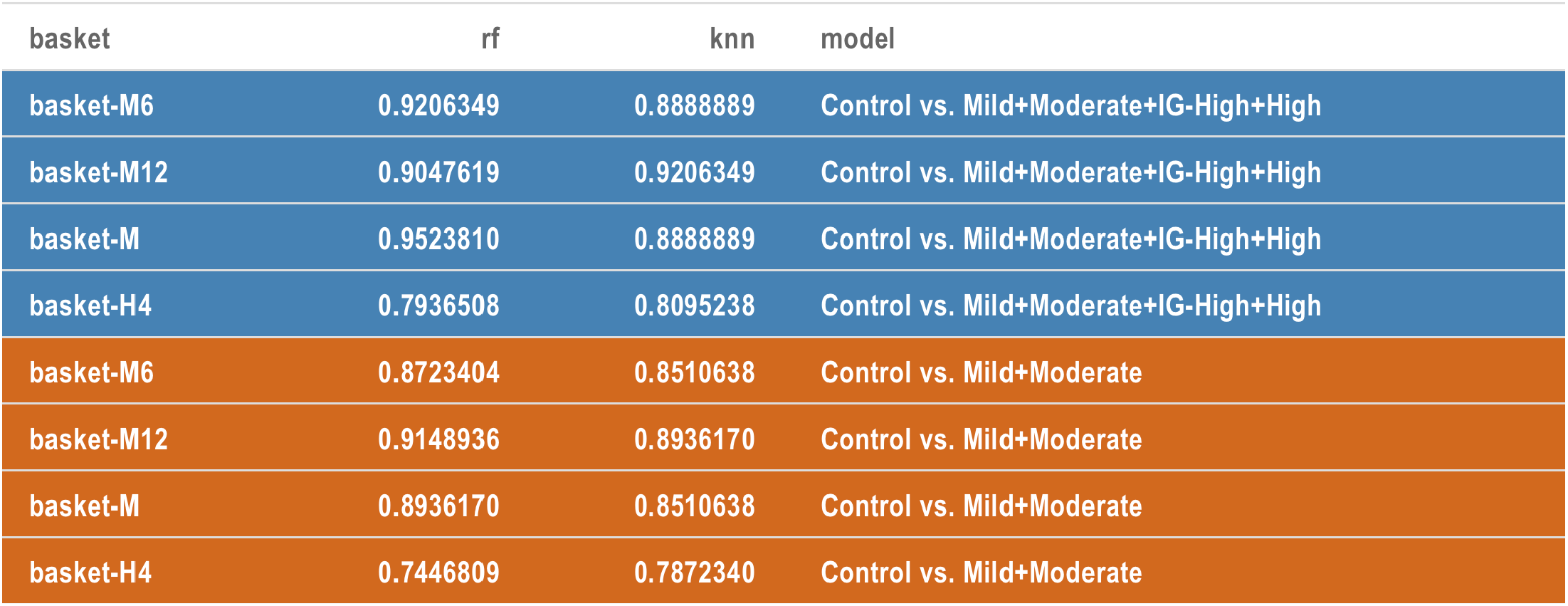
Classification performance of DUX4 baskets by random forest (rf) and KNN algorithm and leave-one-out validation. The orange rows show that when using the longitudinal study as the training model (Control vs. Mild+Moderate), the DUX4 baskets (M6, M12, and M of 34 genes) outperformed the previous used DUX4 robust genes (LEUTX, PRAMEF2, TRAM43, KHDC1L) with above 87% vs. 74% accuracy.

**Supplemental Figure S3.**
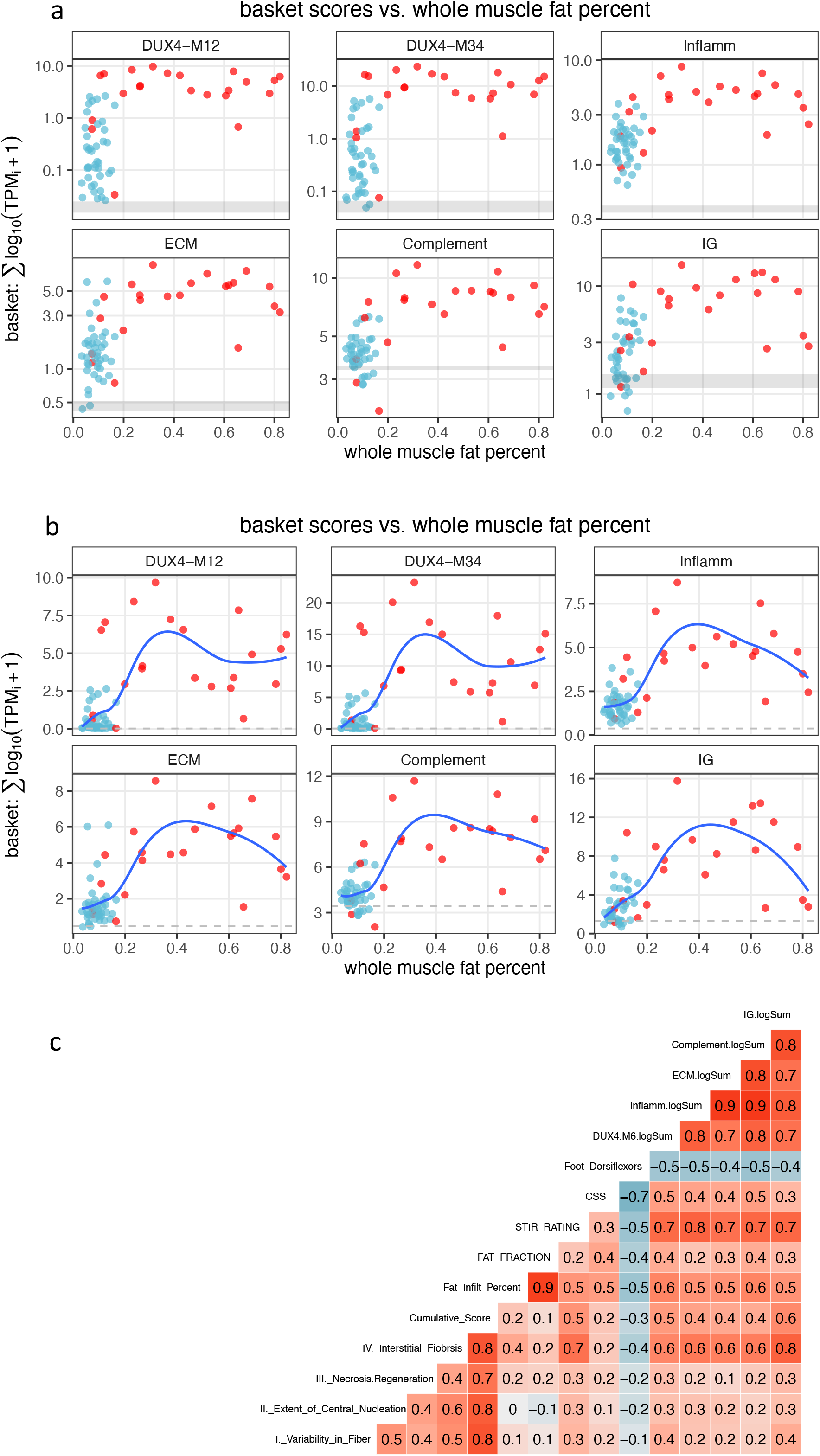
The relationship of whole muscle fat percent to the DUX4, Inflamm, ECM, complement and IG signatures. (a) A scatter plot of the basket scores and the whole muscle fat percent in which the y-axis coordinate is set in log10 scale. The shaded areas present the 95% CI of the historical controls. (b) Same as (a) with the y-axis coordinate set in the original scale. The blue lines present the non-linear (loess) regression. The scores reach highest levels between 0.2 and 0.4 whole muscle fat percent. (c) The comprehensive correlation between the sub-category of the histopathological scores, CSS, MRI characteristics, and basket scores.

**Supplemental Figure S4.**
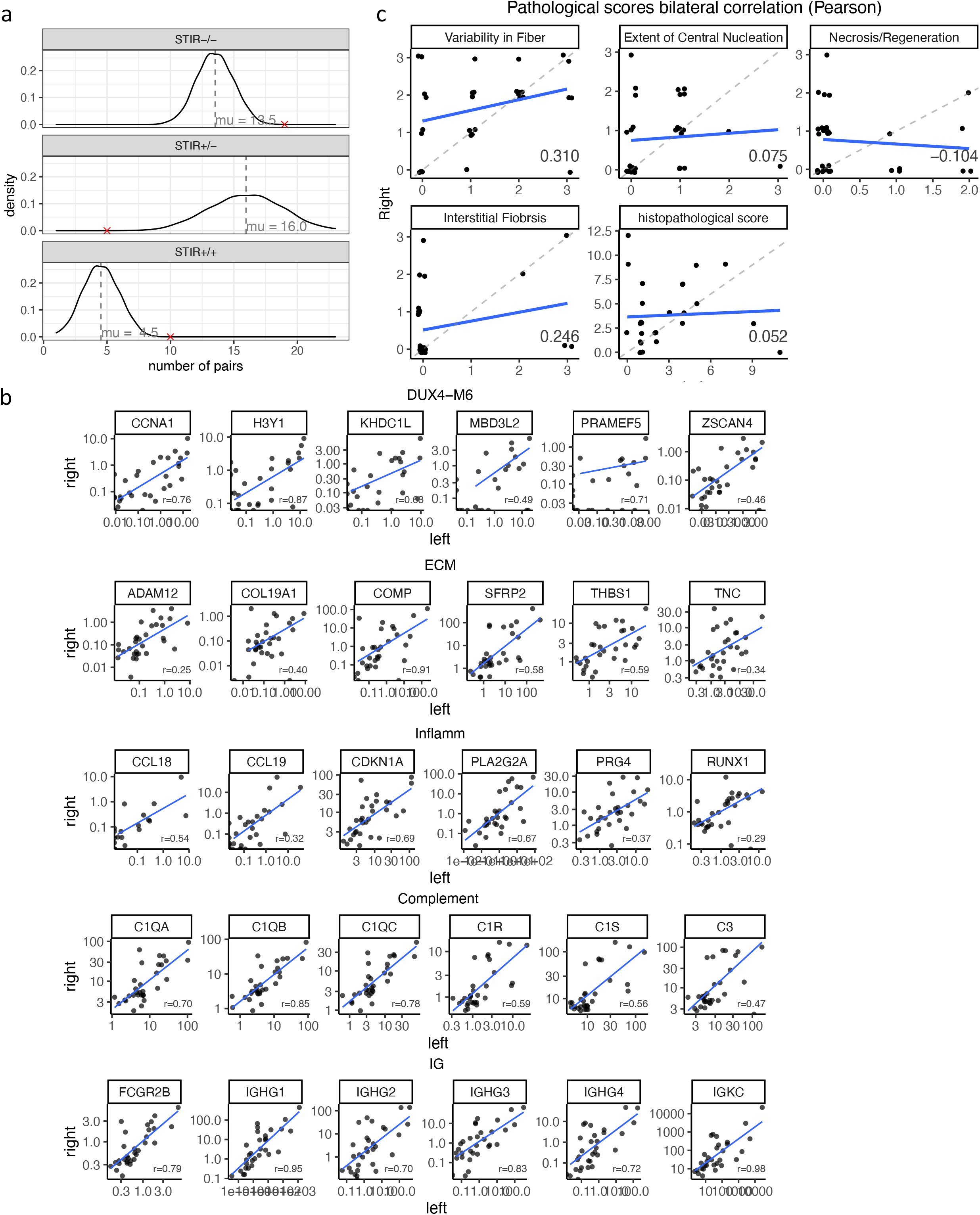
Symmetric trends in left and right tibialis anterior biopsies. (a) Random process simulation showing the distribution of number of STIR-, STIR+ pairs and STIR-/+ discordant pairs. The red marks present the observed number of pairs and the dashed-vertical lines are the expected number of pairs. (b) Scatter plots for gene expression levels (TPM, displayed in log10 scale) between left and right muscle for each gene in the basket. The Pearson correlation was calculated based on TPM. (c) Pathological, MRI and muscle strength correlation between left and right muscle.

